# 4-oxo-2-nonenal Adducts In HDL Are Elevated In Familial Hypercholesterolemia: Identification Of Modified Sites And Functional Consequences

**DOI:** 10.1101/635458

**Authors:** Linda S. May-Zhang, Valery Yermalitsky, John T. Melchior, Jamie Morris, Keri A. Tallman, Mark S. Borja, Tiffany Pleasent, Amarnath Venkataraman, Patricia G. Yancey, W. Sean Davidson, MacRae F. Linton, Sean S. Davies

## Abstract

The lipid aldehyde 4-oxo-2-nonenal (ONE) derived from peroxidation of n-6 polyunsaturated fatty acids and generated in parallel to 4-hydroxynonenal (HNE) is a highly reactive protein crosslinker. Crosslinking of proteins in high-density lipoprotein (HDL) by lipid peroxidation products causes HDL dysfunction and contributes to atherogenesis. While HNE is relatively well studied, the relevance of ONE in atherosclerosis and in modifying HDL has not been examined. In the present study, we found a significant increase in ONE-ketoamide (lysine) adducts in HDL derived from patients with familial hypercholesterolemia (FH) (1620 ± 985.4 pmol/mg) compared to healthy controls (664 ± 219.5 pmol/mg). ONE crosslinked apoA-I on HDL at a concentration of >3 mol ONE per 10 mol apoA-I (0.3 eq), which is 100-fold lower than HNE but comparable to the potent protein crosslinker, isolevuglandin. ONE-modified HDL partially inhibited the ability of HDL to protect against LPS-induced TNFα and IL-1β mRNA expression in murine macrophages. At 3 eq., ONE dramatically decreased the ability of apoA-I to exchange from HDL, from ~46.5% to only ~18.4% (P<0.001). Surprisingly, ONE-modification of HDL or apoA-I did not alter macrophage cholesterol efflux capacity. LC/MS/MS analysis showed modification of Lys12, Lys23, Lys96, and Lys226 of apoA-I by ONE-ketoamide adducts. Compared to other dicarbonyl scavengers, pentylpyridoxamine (PPM) was most efficacious at blocking ONE-induced protein crosslinking in HDL. Our studies show that ONE HDL adducts are elevated in FH who have severe hypercholesterolemia and atherosclerosis and causes HDL dysfunction. We demonstrate the use of PPM in preferentially scavenging ONE in biological systems.

---

Oxidative stress and increased net production of free radicals represent an important pathogenic mechanism in atherosclerosis. Free radicals react with unsaturated fatty acids in a chain reaction to yield lipid peroxides and secondary aldehyde products. These lipid aldehydes are highly reactive and selectively modify proteins or lipids to cause cellular and tissue damage. An important role for reactive aldehydes in the pathogenesis of atherosclerosis is suggested by increases aldehyde-protein adducts in plasma (1,2), aortic atherosclerotic lesions (3–6) and lipoproteins. Adduction to apolipoprotein B in low-density lipoprotein (LDL) enhances their recognition and uptake by macrophages (7,8) which converts macrophages to lipid-laden foam cells (9–11). HDL normally protects LDL from aldehyde adduction by serving as a “sink” for lipid peroxides and their reactive byproducts (12) but, when HDL becomes modified, it results in numerous dysfunctions of HDL (13) (14).

HNE is one of the most investigated aldehydic end-products of oxidative breakdown of membrane n-6 polyunsaturated fatty acids. Its adduction to LDL accelerates its uptake by macrophages (15), and its adduction to HDL causes crosslinks of apoA-I and inhibits its ability to activate lecithin-cholesterol acyltransferase (LCAT) (14). However, the role of its more reactive 4-keto counterpart ONE in atherogenesis is not well studied. Studies characterizing the side-chain modifying chemistry of ONE showed that ONE is a more reactive protein modifier and crosslinking agent than HNE (16–19). ONE is generated in parallel to HNE through the intermediate 4-hydroperoxy-2-nonenal (HPNE) and is a direct product of lipid oxidation (20,21) (**Figure 1**). Like HNE, ONE reacts rapidly with the side chains of Cys, His, Lys residues in proteins via Michael addition, but unlike HNE, ONE also reacts with Arg. Kinetic experiments reveal that the reactivity of ONE towards amino acids of proteins occurs in the following order: Cys >> His > Lys > Arg (26). While both HNE and ONE form Schiff bases with Lys, only ONE is capable of forming the 4-ketoamide adduct (24, 31).

**Figure 1.**
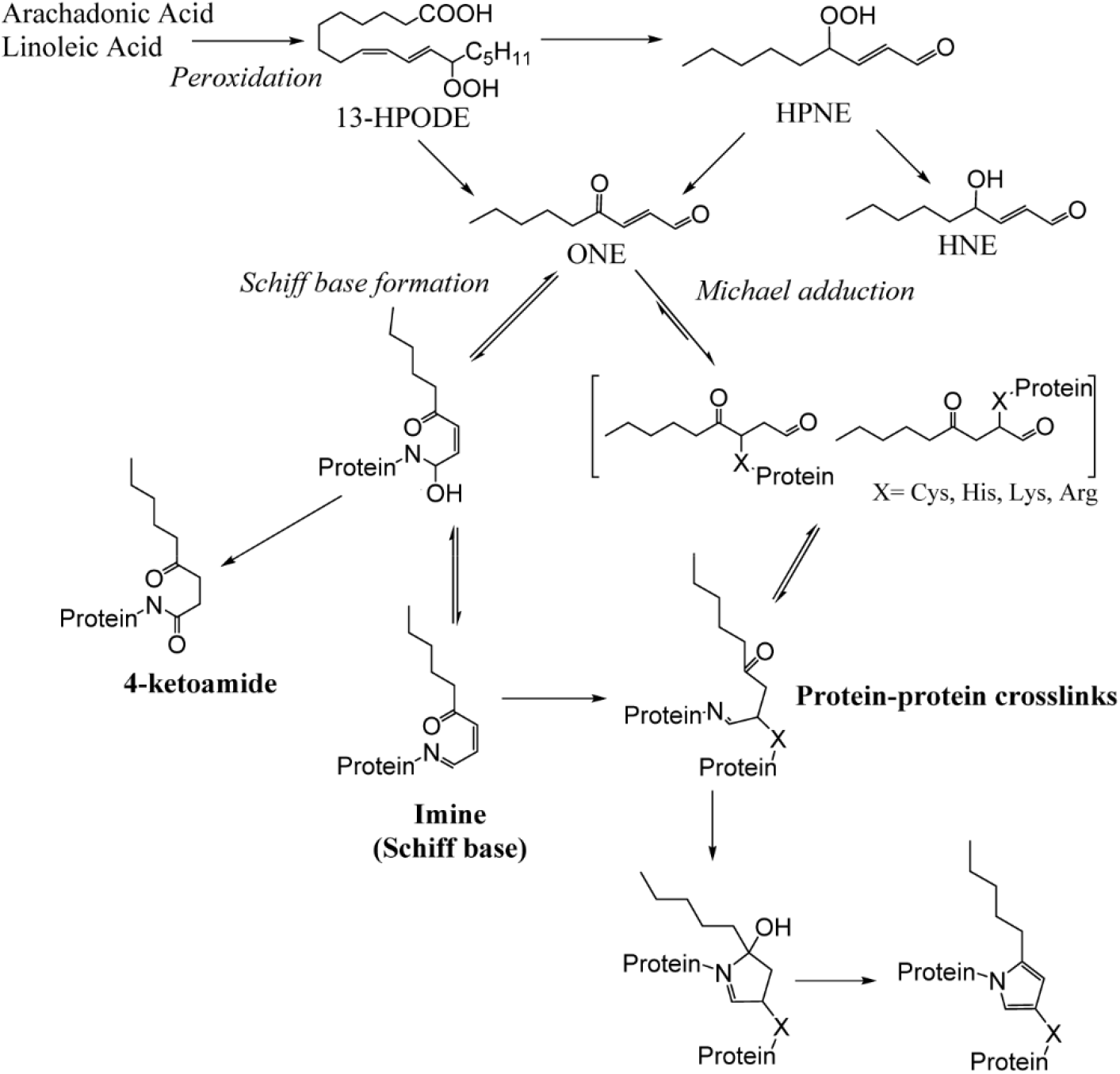
Formation of ONE from the peroxidation of arachidonic or linoleic acid and its adduction to proteins.

In vivo, HNE- and HPNE-specific epitopes exist in atherosclerotic plaques (22,23). Studies using carnosine as an aldehyde scavenger show the presence of carnosine-ONE in oxidized LDL (24). However, no other studies to our knowledge have examined the biological consequences of ONE modification of lipoproteins in atherosclerosis. In this study, we utilized LC/MS/MS to measure ONE-lysine adducts in HDL derived from patients with familial hypercholesterolemia compared to control healthy age-matched subjects. We also identified the amino acid sites of apoA-I that ONE targets and determined the consequences of ONE modification on HDL function. We demonstrate for the first time that ONE-HDL adducts are elevated in atherosclerosis. Interestingly, ONE causes HDL dysfunction in terms of rendering HDL unable to protect against LPS-induced macrophage activation but does not alter its ability to promote macrophage cholesterol efflux.

## RESULTS

### ONE-lysine adducts in human HDL are elevated in atherosclerosis

Levels of ONE protein adducts in HDL were determined in patients with familial hypercholesterolemia (n=7) compared to healthy control volunteers (n=8). One of the FH patients had homozygous FH and six of the subjects had severe heterozygous FH and were undergoing regular LDL apheresis. From these patients, plasmas were collected before LDL apheresis. Because ONE-induced crosslinking generates multiple chemical structures depending on the microenvironment (25–27), we focused instead on quantifying the lysine 4-ketoamide adduct. These monoadducts are irreversible, highly stable and longer-lived (25). By LC/MS/MS, we found that the levels of ONE-ketoamide adducts were significantly higher (P<0.05) in familial hypercholesterolemia (1620 ± 985.4 pmol/mg protein) than in controls (664 ± 219.5 pmol/mg protein) (**Figure 2**). These results are first to demonstrate that ONE-protein adducts are elevated on human HDL in conditions of severe hypercholesterolemia associated with atherosclerosis.

**Figure 2.**
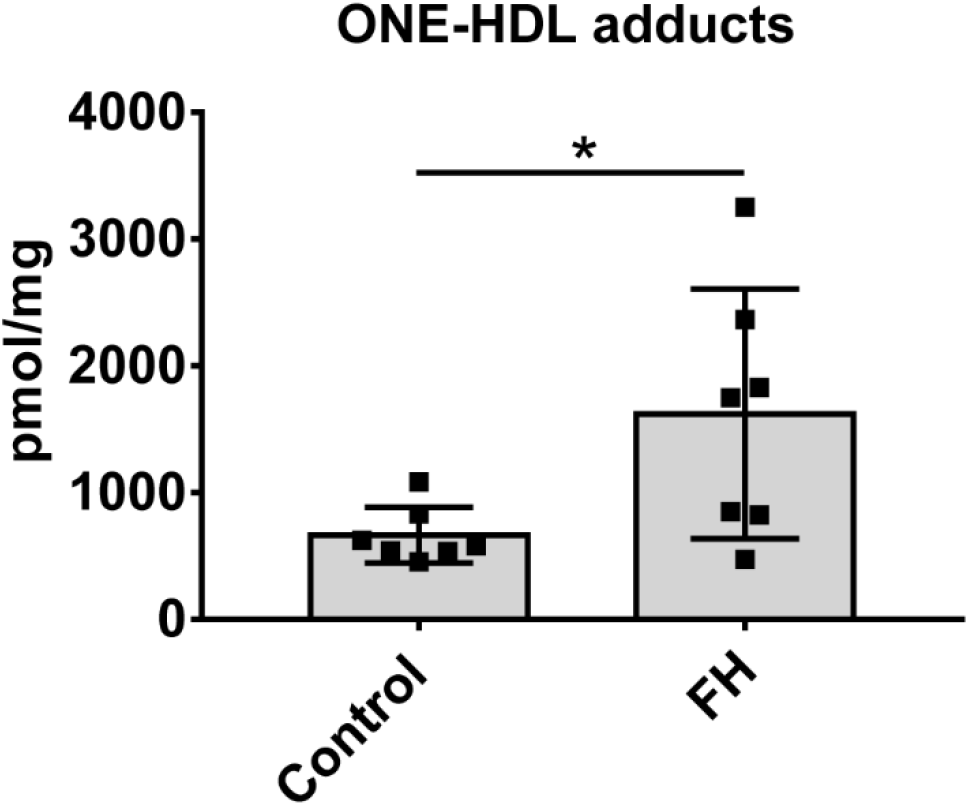
ONE-protein adducts are increased in HDL isolated from plasma of patients with familial hypercholesterolemia (FH). Plasma was isolated from the blood of FH patients (n=7) and healthy volunteers (n=8) and subjected to FPLC to isolate the HDL. Levels of ONE-lysine ketoamide adducts were determined by LC/MS/MS. Values are shown as mean ± SD. Statistical significance was calculated by Student’s t-test. *P<0.05.

### ONE potently crosslinks HDL proteins but does not significantly alter HDL size distribution

For these studies, we first determined the concentration of ONE needed to modify HDL in vitro that would generate the levels of protein adducts within same range as we found in our in vivo samples. Levels of ketoamide adducts were determined in control HDL modified with a range of ONE concentrations (**Supplementary Figure 1**). Approximately 0.3 eq. ONE (1 ONE per 3 apoA-I molecules) to modify HDL yielded ketoamide adduct levels (2648 ± 1156 pmol/mg) similar to levels measured from circulating HDL in FH patients. Thus, for our biological assays, we used a range from 0.1 to 3 eq. ONE to modify HDL so as to span the levels found in circulating HDL up to theoretical conditions where ONE may be produced more locally and thus concentrated such as within the arterial wall.

In addition to forming ketoamide monoadducts, ONE potently crosslinks HDL proteins starting at only ~0.1 eq (**Figure 3A**) as indicated by SDS-PAGE and Coomassie Blue staining to visualize the proteins. Structural proteins apoA-I and apoA-II are crosslinked beginning at 0.1 eq (**Figures 3B, C**). This concentration needed to crosslink is approximately equal to that needed for isolevuglandins (a family of 4-ketoaldehydes) and 10-fold lower than HNE (28). At 10 eq., the monomers of apoA-I and apoA-II begin to disappear, as they are crosslinked and form high molecular weight oligomers. The patterns of these high molecular weight oligomers appear different from that formed by isolevuglandin-crosslinking (28), indicating different reaction chemistries between the two dicarbonyls. Interestingly, 3 eq. ONE was required to crosslink apoA-IV (**Figure 3D**), another HDL protein that can activate LCAT (29) and has antioxidant properties (30,31). Potentially, ONE targets apoA-I and apoA-II due to their greater abundance on HDL. The data indicates that ONE is a very reactive electrophile that potently crosslinks HDL proteins, making ONE equally as reactive as the isolevuglandins.

**Figure 3.**
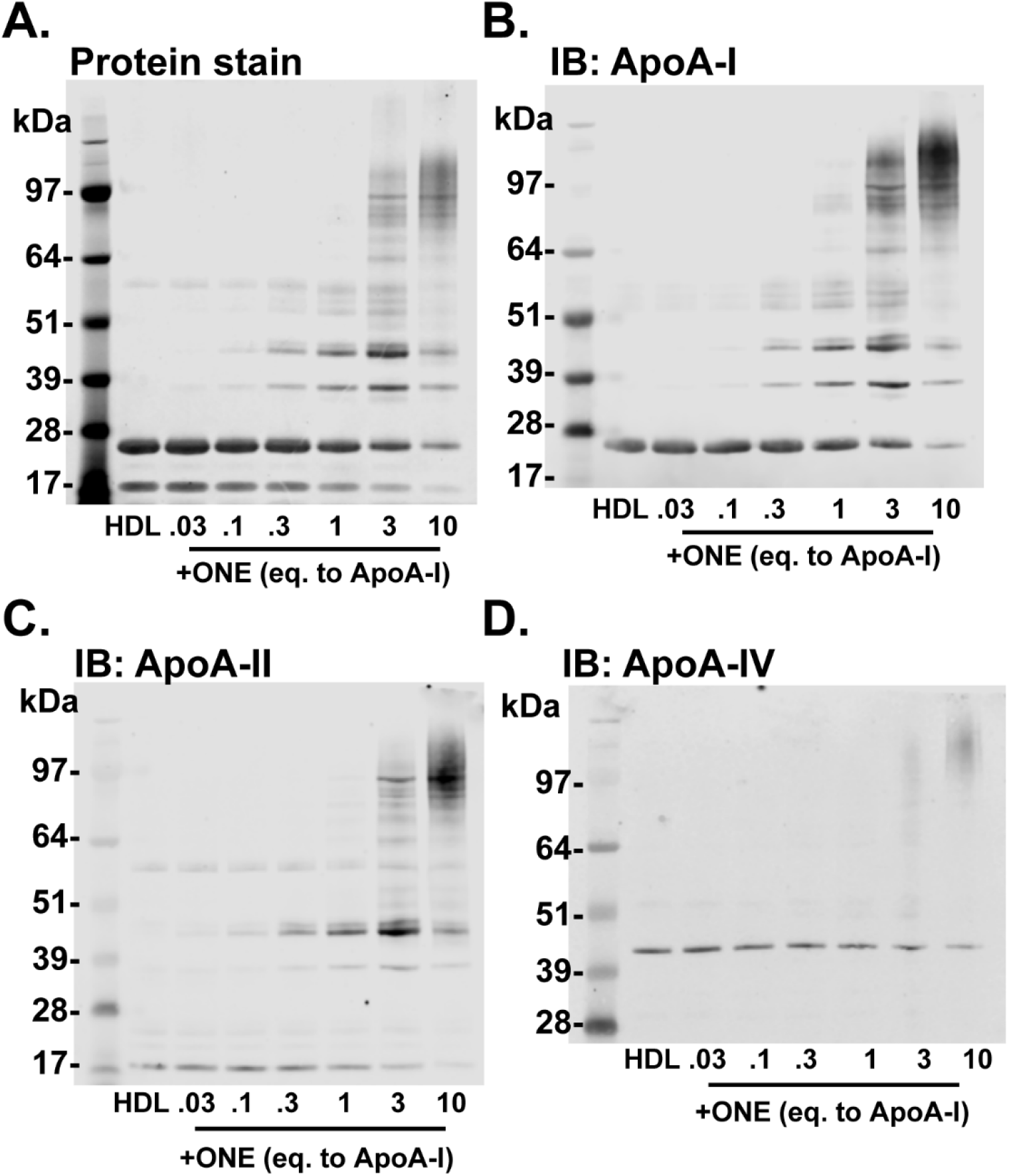
ONE crosslinks HDL proteins beginning at 0.1 molar eq. to apoA-I. Crosslinking of HDL proteins by ONE was determined by (A) Coomassie blue staining and (B-D) immunoblotting (IB) against apoA-I, apoA-II, and apoA-IV. HDL isolated from normal healthy subjects by density gradient ultracentrifugation was subjected to *ex vivo* modification of ONE. SDS-PAGE and Western blots are representative of experiments performed three times.

Because isolevuglandins crosslink HDL to produce a HDL subpopulation of larger size (28), we sought to determine the impact of ONE modification on HDL size distribution. Using fast protein liquid chromatography (FPLC), we found that both unmodified and ONE-modified HDL fractionate into two main subpopulations: spherical and lipid-poor HDL (**Figure 4A**). However, ONE did not significantly change the size or distribution of HDL, suggesting that the particles were not fusing to form larger sized HDL, unlike what it seen with isolevuglandin modification. Native PAGE gel electrophoresis of reconstituted HDL modified by ONE also shows no change in electrophoretic mobility of these particles (**Figure 4B**). Together, the data would suggest that ONE mainly forms intramolecular crosslinks or crosslinks of proteins originally on the same HDL particles, rather than crosslinking proteins on adjacent particles that leads to fusion of the HDL particles.

**Figure 4.**
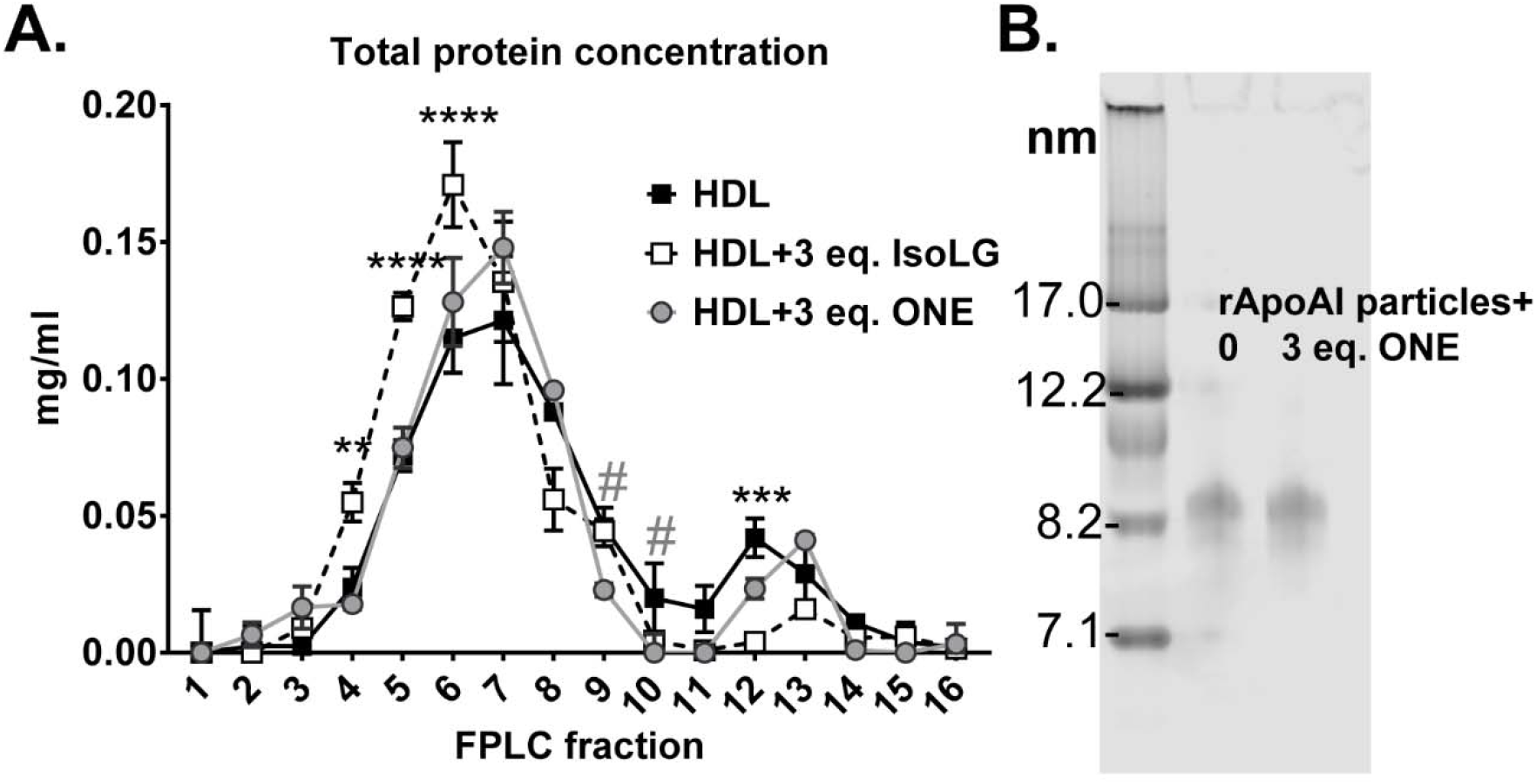
ONE does not significantly alter HDL size or distribution as indicated by A) size exclusion chromatography and by B) native PAGE gel electrophoresis. HDL was subjected to ex vivo modification of ONE under aqueous conditions 37°C overnight. HDL was subjected to FPLC using the Superdex 200 column using A280 monitoring. Protein concentration of fractions was assessed using the Bradford assay. Values are represented as mean ± SD. Statistical significance was calculated by 2-way ANOVA with Dunnett’s multiple comparisons test compared to HDL. *represents statistical significance between HDL and IsoLG-HDL, while # represents significance between HDL and ONE-HDL. #P<0.05, **P<0.01, ***P<0.005, ****P<0.001. Native PAGE gel electrophoresis was run under standard conditions and gel was stained with Coommassie G250 to visualize particles.

### ONE-modified HDL have lower HDL -apoA-I exchange but no effect on total macrophage cholesterol efflux

Previously we showed that HDL highly crosslinked by isolevuglandins has reduced ability to exchange apoA-I, which correlates with reduced ability to promote macrophage cholesterol efflux (28). Because ONE potently crosslinks HDL proteins, we sought to characterize HDL-apoA-I exchange in ONE-modified HDL. Using the method by Borja et al, which utilizes electron paramagnetic resonance (EPR) to detect conformational changes resulting from lipid binding of nitroxide-labeled apoA-I to HDL (32), we found that ONE modification of HDL dose-dependently reduced apoA-I exchangeability. While unmodified HDL had an apoA-I exchange rate of 46.5±5.6%, HDL exposed to 1 eq. ONE yielded only 19.8±5.7% exchange (P<0.01) (**Figure 5**).

**Figure 5.**
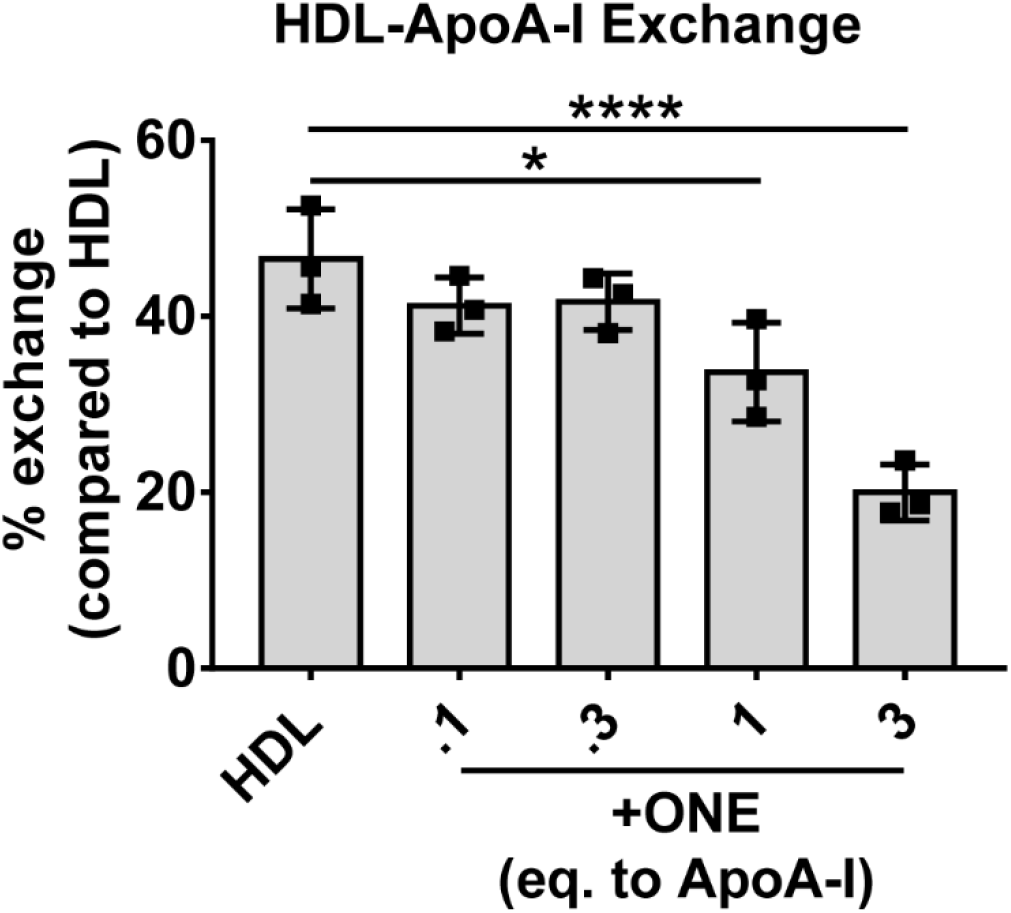
Modification of HDL by ONE inhibits the exchangeability of apoA-I on HDL. HDL-ApoA-I exchange was analyzed by electron paramagnetic resonance as described in Methods. Reactions were performed at a constant apoA-I concentration of 1 mg/ml. Samples were assayed in triplicate. Results are plotted as mean ± SD. Statistical significance was calculated by one-way ANOVA with Dunnett’s multiple comparisons compared to unmodified HDL (control). *P<0.05; ****P<0.0001.

A decrease in apoA-I exchange often results in less efficient cholesterol mobilization from macrophages via ABCA1, so we examined effect of ONE modification on cholesterol efflux as previously described (28). Using peritoneal murine macrophages isolated from apoE^-/-^ mice (to exclude the effect of macrophage apoE) and loaded with acetylated LDL, we found that ONE modification did not affect the ability of HDL to efflux ^3^H-cholesterol even at concentrations up to 3 eq. (**Figure 6**). This showed that although apoA-I exchange was dramatically affected by ONE, the capacity to efflux cholesterol was unaffected.

**Figure 6.**
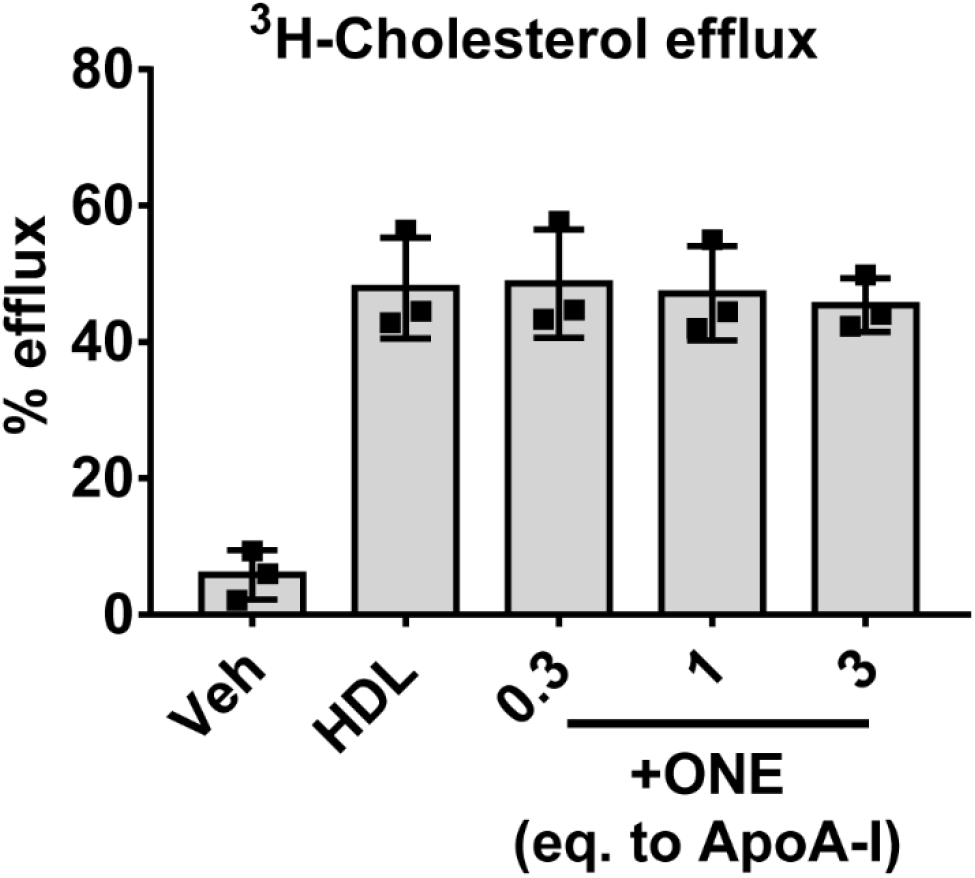
Modification of HDL by ONE does not affect cholesterol efflux from apoE deficient murine macrophages. HDL isolated from normal healthy subjects by DGUC was subjected to ex vivo modification of ONE. Macrophage cholesterol efflux was assessed using thioglycollate-induced macrophages harvested from the peritoneum of apoE deficient mice, and loaded with ^3^H cholesterol and acetylated LDL. Vehicle denotes cell culture media with no HDL added to the cells. Efflux of ^3^H to unmodified and modified HDL was calculated based on radioactive counts in the supernatant after 4 h and normalized to HDL control. Results from three individual experiments with replicate wells per treatment are plotted as mean ± SD.

### ONE modification negates the ability to protect against LPS-stimulated macrophage inflammatory response

We next determined if ONE modification rendered HDL dysfunctional in protecting against LPS-induced proinflammatory cytokine expression in macrophages. Previously, we found that isolevuglandin-modified HDL not only rendered HDL ineffective at preventing LPS-induced inflammatory response but induced a greater response than LPS alone (28). In the current study, we found that while unmodified HDL inhibited the LPS-induced expression of *Tnfα, IL-1β*, and *IL-6* by 77.0±2.3%, 59.9±2.7%, 37.3±12.8% respectively, modification with 0.1 eq. ONE ablated its ability to prevent *Tnfα, IL-1 β, and IL-6* expression, resulting in 94.1 ± 20.9%, 108.9 ± 28.0%, 97.2± 26.5% of LPS respectively at 3 eq. ONE (**Figure 7**). However, these levels of cytokine expression are similar to LPS induction alone, indicating that ONE-modified HDL do not potentiate a greater inflammatory response by these macrophages.

**Figure 7.**
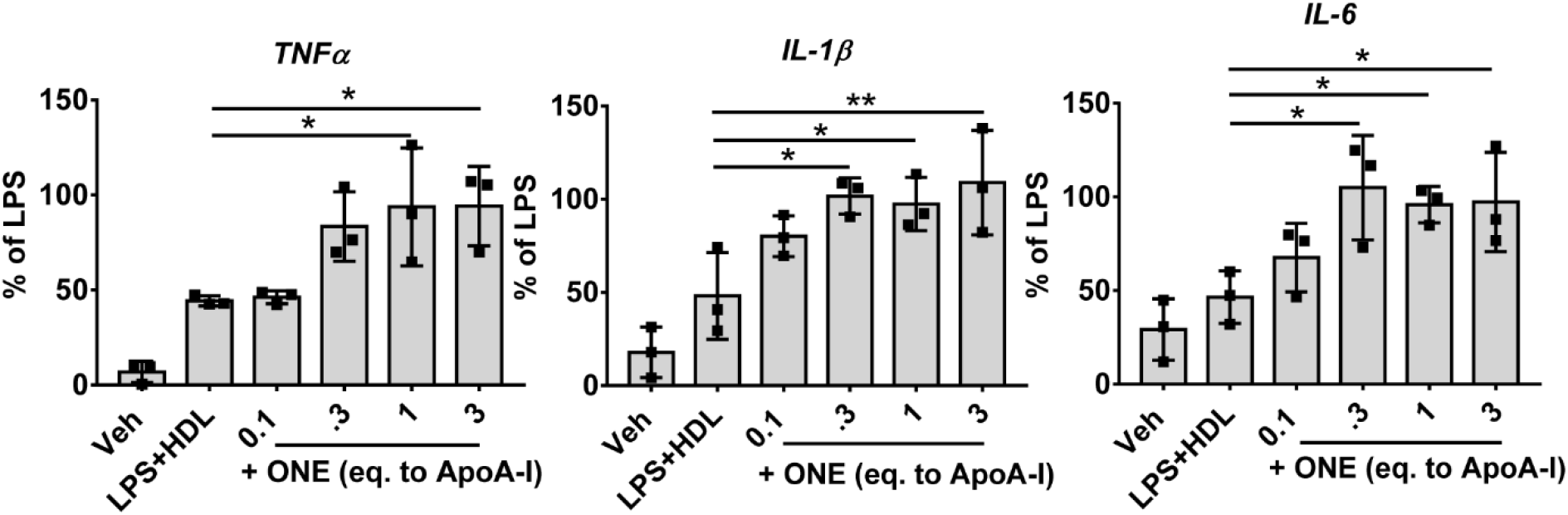
Modification of HDL by ONE renders HDL no longer protective against LPS-stimulated inflammatory cytokine expression. HDL isolated from normal healthy subjects by DGUC was subjected to ex vivo modification of ONE. Thioglycollate-elicited peritoneal macrophages were treated with LPS along with ONE modified HDL for 4 hours. Experiments were performed independently three times with three wells per treatment. Results from three individual experiments with 3-4 wells per treatment are plotted as mean ± SD. Statistical significance was determined by one-way ANOVA with Dunnett’s multiple comparisons compared to LPS+HDL (control). *P<0.05; **P<0.01.

### LC/MS/MS identification of ONE modified lysine residues in apoA-I

To identify the amino acid targets of ONE in apoA-I, we used reconstituted (r)HDL generated using recombinant human ApoA-I, phosphatidylcholine, and free cholesterol. rHDL particles represent simplified systems that have been used for decades to study the HDL-like lipid bound state of apoA-I. rHDL was modified with 3 eq. ONE overnight. After reaction quenching, the particles were delipidated by a chloroform/methanol extraction and the resulting protein was exhaustively digested with trypsin. The resulting peptides were analyzed by LC/MS/MS and the data was interrogated for known mass adducts resulting from ONE chemical modifications. A representative MS/MS identification of the apoA-I peptide AKVQPYLDDFQK with a ONE ketoamide adduct on Lys 97 is shown in **Figure 8**. The spectrum shows the MS/MS spectrum with b+ and y+ series ions highlighted in red and labeled in blue. The table shows the identified ions bolded in red. **Table 1** summarizes all the identified peptides and ONE adducts resulting from three independent experiments. The identified residues include ketoamide adducts (+154.0994 amu) on Lys12, 23, 96, and 226 in human apoA-I in triplicate experiments. One of the three experiments also identified ketoamide adducts on Lys 94 and 118, while another experiment identified Michael adducts (+156.1200 amu) on Lys 133 and 226.

**Figure 8.**
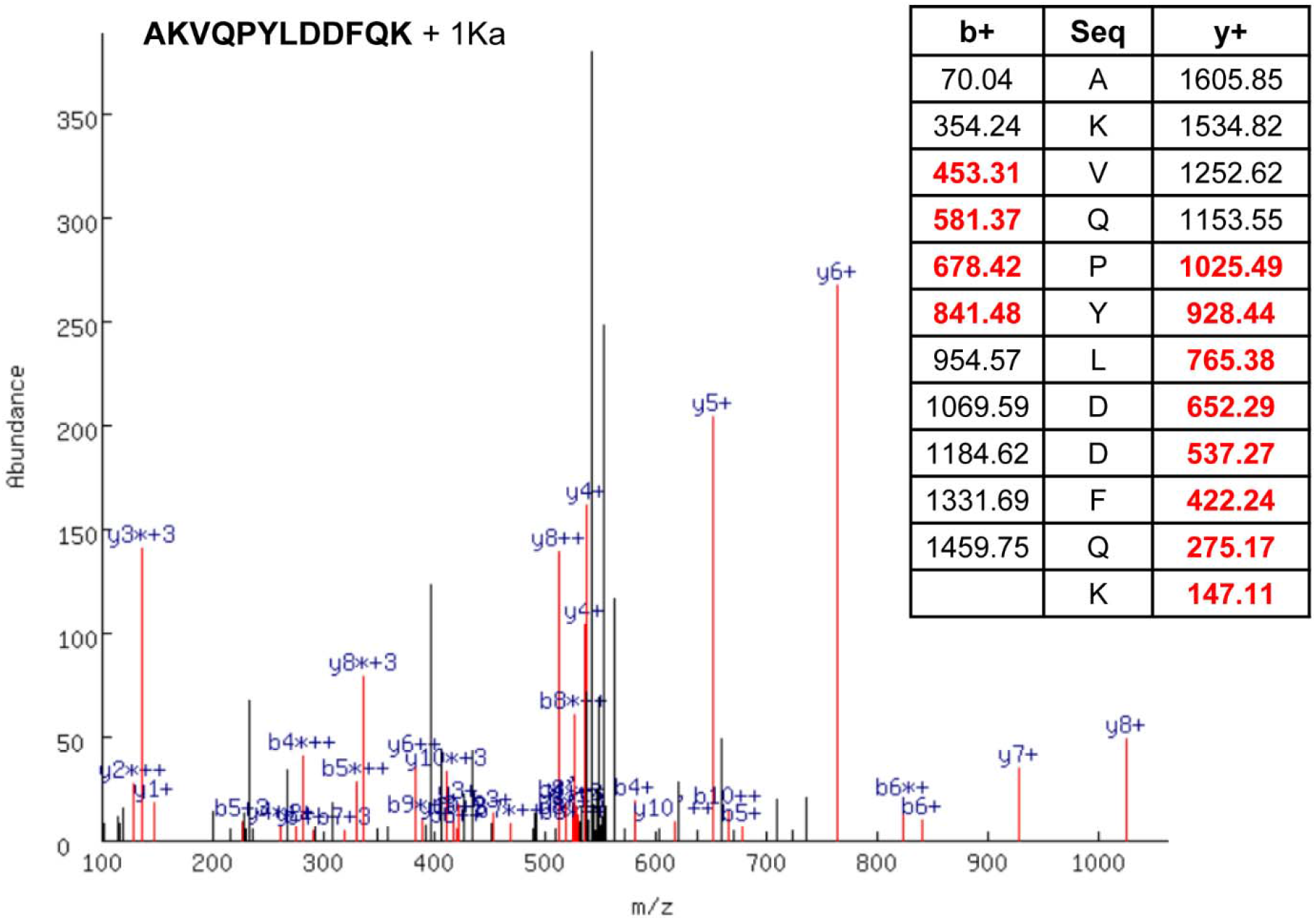
Representative MS/MS identification of the apoA-I peptide AKVQPYLDDFQK with a ONE ketoamide adduct on Lys 97. The spectrum shows the MS/MS spectrum with b+ and y+ series ions highlighted in red and labeled in blue. The table shows the identified ions bolded in red.

**Table 1.**
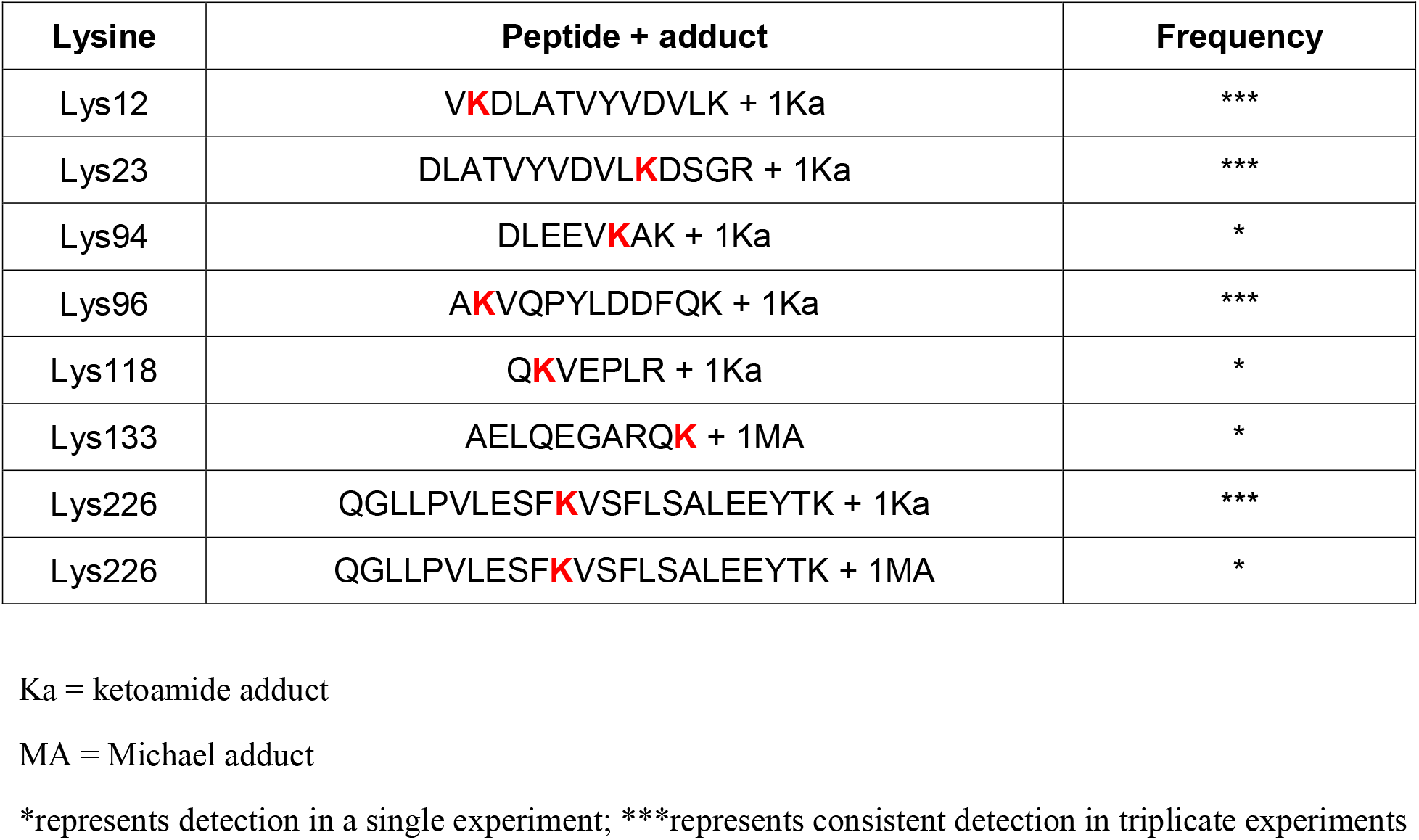
Summary of all the identified peptides and ONE adducts resulting from three independent experiments. The identified residues include ketoamide adducts (+154.0994 amu) on Lys12, 23, 96, and 226 in human apoA-I in triplicate experiments. One of the three experiments also identified ketoamide adducts on Lys 94 and 118, while another experiment identified Michael adducts (+156.1200 amu) on Lys 133 and 226.

| Lysine | Peptide + adduct | Frequency |
| --- | --- | --- |
| Lys12 | V~~K~~DLATVYVDVLK + 1Ka | *** |
| Lys23 | DLATVYVDVL~~K~~DSGR + 1Ka | *** |
| Lys94 | DLEEV~~K~~AK + 1Ka | * |
| Lys96 | A~~K~~VQPYLDDFQK + 1Ka | *** |
| Lys118 | Q~~K~~VEPLR + 1Ka | * |
| Lys133 | AELQEGARQ~~K~~ + 1MA | * |
| Lys226 | QGLLPVLESF~~K~~VSFLSALEEYTK + 1Ka | *** |
| Lys226 | QGLLPVLESF~~K~~VSFLSALEEYTK + 1MA | * |

Ka = ketoamide adduct
MA = Michael adduct
\*represents detection in a single experiment; \*\*\*represents consistent detection in triplicate experiments

### Pentylpyridoxamine is more effective at scavenging ONE than other scavengers

2-aminomethylphenols have been shown to scavenge ONE *in situ*, with PPM and chloro-salicylamine possessing the fastest second-order reaction rate (33). However, unknown is the effectiveness of these scavengers in preventing ONE from reacting with proteins embedded in a lipophilic environment of the HDL surface. Incubating the scavengers and ONE together with HDL overnight and assessing protein crosslinking by SDS-PAGE, we found that PPM was significantly more effective at scavenging ONE compared to salicylamine (2-hydroxybenzylamine) or its analogues (fluoro-salicylamine, F-SAM; chloro-salicylamine, Cl-SAM) (**Figures 9a, b**). Furthermore, PPM at 10 eq. was able to prevent 1 eq. ONE-induced HDL dysfunction in protecting against LPS-stimulated Tnfα expression in macrophages while its inactive analogue, pentylpyridoxine (PPO), did not. This demonstrates the effectiveness of PPM in preventing ONE-induced crosslinking as well as HDL dysfunction in a biological system.

**Figure 9.**
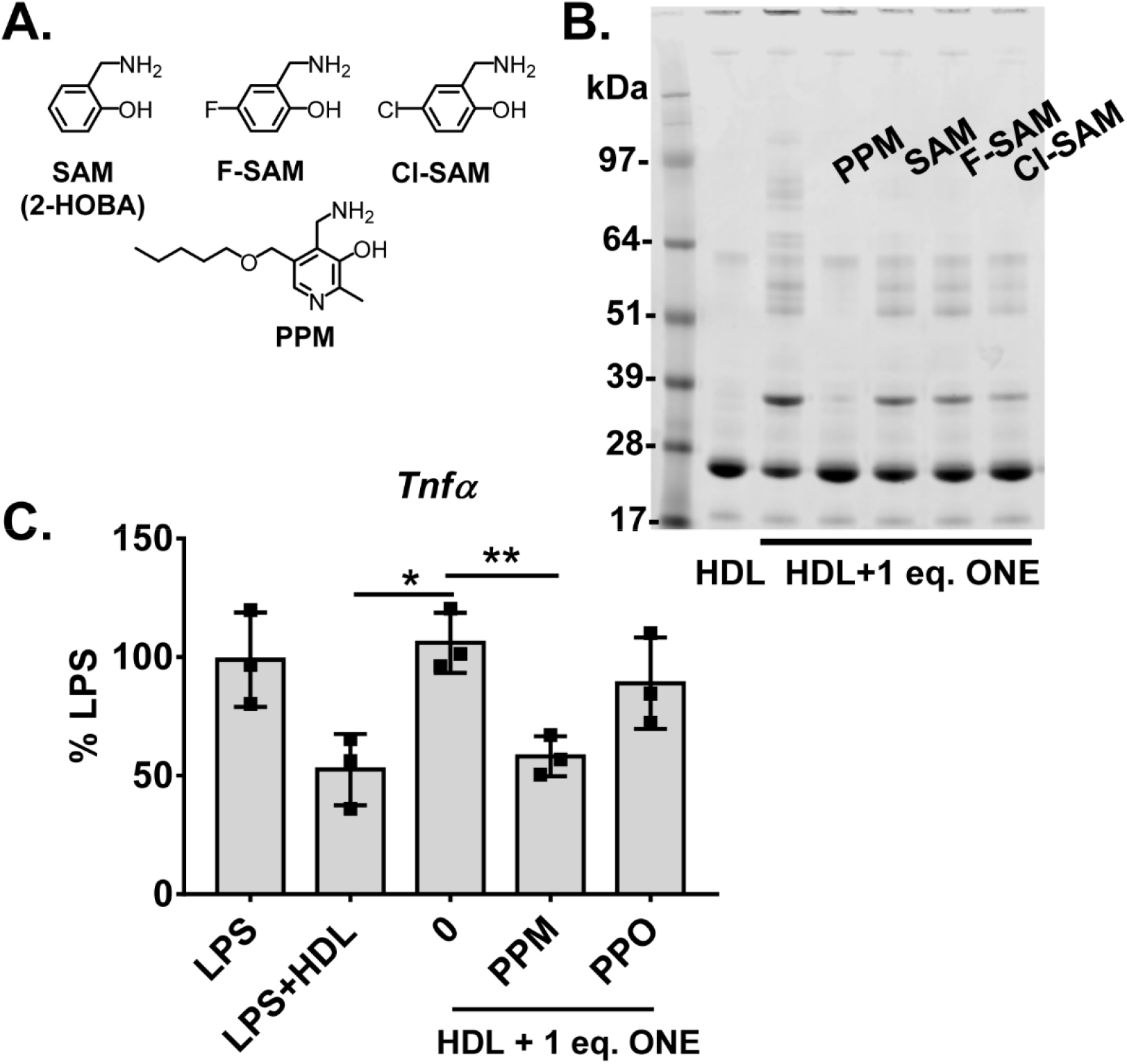
Effect of salicylamine analogues in scavenging ONE. (A) Molecular structures of salicylamine and various analogues (fluoro-salicylamine, F-SAM; chloro-salicylamine, Cl-SAM; pentylpyridoxamine, PPM). (B) Effect of analogues in preventing ONE-induced HDL protein crosslinking as shown by Commassie G250 stained SDS-PAGE gel of unmodified and 1 eq. ONE-modified HDL. (C) Demonstration of PPM (10 eq) preventing ONE-induced HDL dysfunction in protecting against LPS-stimulated Tnfα expression in thioglycollate-elicited peritoneal macrophages while its inactive analogue, PPO, did not. Statistical significance was determined by one-way ANOVA with Dunnett’s multiple comparisons compared to LPS+1 eq. ONE+HDL. Results from three individual experiments with 3 wells per treatment are plotted as mean ± SD. *P<0.05; **P<0.01.

## DISCUSSION

Growing evidence suggests that modification of HDL by reactive lipid aldehydes renders HDL dysfunctional which contributes to the pathogenesis of atherosclerotic disease. While ONE is far more reactive than HNE, its role in atherosclerosis is unknown. In the present study, we show that ONE-protein adducts are elevated in human atherosclerosis. ONE crosslinks HDL proteins at low molar concentrations but does not alter HDL size. While ONE-modification alters HDL function in certain aspects (such as HDL apoA-I exchange or anti-inflammation), it does not affect macrophage cholesterol efflux. We also identify 4 lysines that ONE consistently targets in apoA-I. Finally, we show that the dicarbonyl scavenger, PPM, is most effective at scavenging ONE from crosslinking HDL proteins.

Since the 4-ketoamide adducts are known as the major long-lived protein adducts, we initially assayed for the ONE-ketoamide adducts in patient HDL. We found a significant elevation in ONE-ketoamide adducts in the HDLs of patients with familial hypercholesterolemia, a level which can be achieved by incubating 0.3 molar eq. of ONE (to apoA-I) with native HDL. Thus, for our *in vitro* experiments, we used a 10-fold range around 0.3 eq to represent modified HDL derived from the steady-state plasma to a localized environment such as within an atherosclerotic lesion where oxidants are generated.

While ONE is structurally analogous to HNE, the ketone at the C4 position makes ONE much more reactive towards protein nucleophiles (16,34). We demonstrate that ONE crosslinks the main structural proteins of HDL, apoA-I and apoA-II, at very low concentrations (0.1-0.3 eq to apoA-I.), making ONE just as reactive as the potent isolevuglandins in protein crosslinking (28). Modification of HDL by above 1 eq. ONE produces multimers of apoA-I and apoA-II that migrate as distinct oligomers through an SDS gel, which contrasts with the ‘smear’ of oligomers created by isolevuglandin-induced crosslinking. This suggests that the proteins crosslinked by ONE differ to some extent from those crosslinked by isolevuglandins, which may contribute to differences in the dysfunctions of HDL induced by these two reactive dicarbonyls.

The exchangeability of apoA-I in HDL governs many of HDL’s functions, such as its ability to promote cholesterol efflux via ABCA1 (35). Decreases in HDL-apoA-I exchange were observed in both animal models of atherosclerosis and in human atherosclerosis (32). Decreases in HDL-apoA-I exchange also correlated with decreased cholesterol efflux capacity of isolevuglandin-modified HDL (28). When we measured the rate of exchange between spin-labeled apoA-I and ONE-modified HDL, we found a decrease in apoA-I exchangeability that depended on extent of ONE modification. This dose-dependent decrease is comparable to that of isolevuglandin-modified HDL. Crosslinking of HDL proteins may hinder the ability of HDL-bound apoA-I to exchange with exogenously added lipid-free apoA-I associated with the particles, thus resulting in decreased HDL-apoA-I exchange.

While HDL-apoA-I exchange correlates with cholesterol efflux capacity, we find that ONE-modification of HDL or apoA-I did not alter the ability of HDL to induce cholesterol efflux from macrophages at any of the tested concentrations. This finding was initially surprising because ONE dramatically decreased HDL-apoA-I exchange as well as significantly crosslinked HDL proteins. Crosslinking of HDL proteins by isolevuglandins correlated with the significant reduction of its cholesterol efflux capacity (28). However, size exclusion separation of spherical and lipid-poor HDL shows that ONE-modification does not appear to alter HDL size or distribution of spherical versus lipid poor particles. This contrasts with isolevuglandin modification, which increases HDL size (28) and decreases the population of lipid-poor particles (*data not shown*). Since small lipid poor HDL more efficiently promotes cholesterol efflux than larger spherical HDL via ABCA1 (36), it may be possible that ONE modification does not impact cholesterol efflux due to the lack of alteration of HDL size and population of the small particles. Additionally, not all endogenous crosslinkers of HDL proteins impair function. Peroxidase-generated tyrosyl radicals crosslinks apoA-I and apoA-II in HDL but enhances its ability to promote cholesterol efflux (37,38).

Another function of HDL is to protect against inflammation partially by neutralizing LPS (39–42). Previously, we found that modification of HDL with isolevuglandins not only completely blocks the ability of HDL to inhibit LPS-induced cytokine expression in macrophages but also further potentiates *Il-1β* and *Il-6* expression induced by LPS. In this study, we find that ONE modification of HDL blocks its ability to inhibit LPS-induced *Tnfα, Il-1β*, and *Il-6* expression beginning at 0.3 eq. ONE. However, increasing ONE modification does not further potentiate expression of these cytokines by LPS. Of note, modification of HDL with HNE or succinaldehyde does not potentiate LPS expression either (28). These results suggest that the potentiation of cytokine expression induced by isolevuglandin modification of HDL results from a mechanism (e.g. formation of a receptor agonist) distinct from the mechanism whereby HDL inhibits LPS signaling (e.g. neutralization of LPS by apoAI). Potentiation of expression might result from isolevuglandin modification specifically forming receptor agonists while the block of inhibition might result from apoA-I modified by a wider array of aldehydes being unable to bind and neutralize LPS.

Quantitative analysis of proteolytic digests of apoA-I by LC/MS/MS revealed that ONE consistently formed ONE-ketoamide adducts on four lysine residues (Lys12, Lys 23, Lys96, Lys226) in apoA-I. One of the three experiments also identified ketoamide adduct on Lys118, while another experiment identified Michael adducts on Lys 133 and 226. This may suggest that these sites are lower frequency targets. Unlike HNE (13), we saw no evidence of Michael adducts on any histidine residues. Interestingly, when superimposed onto the published crystal structure of an apoA-I dimer that has been reported to bear strong similarities to the lipid-bound form (43), these Lys residues are all highly solvent ex-posed (**Figure 10**). In fact, lysines 96, 118 and 133 are pointing almost directly outward from the double belt structure that likely forms the basis of apoA-I’s encap-sulation of HDL lipids (44). Lys12 was previously shown to be susceptible to glycation and important in apoA-I’s antiinflammatory function (43). Lys23 resides in a region potentially significant in interactions with lipoprotein-binding protein (44), which plays an important role in not only facilitating transfer of LPS to soluble CD14 but also to neutralize these complexes via transfer to HDL (45). Lys96 and Lys133 reside in helix 3 and 5, respectively, and are potentially involved in LCAT activation (46) and also in interactions with lipoprotein-binding protein (44). Our finding that ONE modifies Lys implicated in the interaction with lipoprotein-binding protein suggest that the inability of ONE-modified HDL to inhibit LPS induced macrophage activation likely results from reduced ability to bind and neutralize LPS.

**Figure 10.**
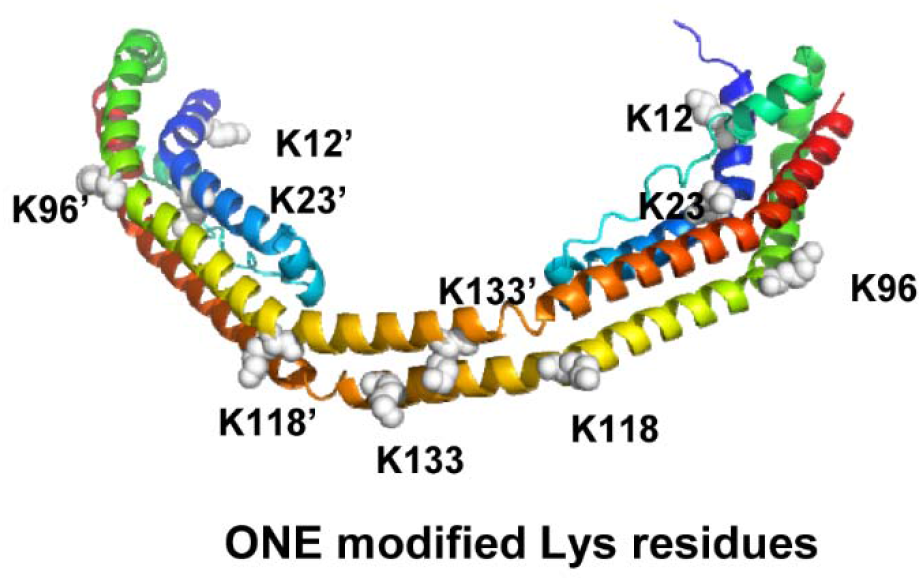
Superimposed ONE targeted lysines onto the published crystal structure of an apoA-I dimer that has been reported to bear strong similarities to the lipid-bound form.

Interestingly, Lys226 was one of the Lys residues found to be highly modified by ONE. Lys226 is also a major target of acrolein (47) and malondialdehyde (MDA) (13). Lys226 appears to be important in cholesterol efflux (48). Deletions of helix 10 where Lys226 resides (49) or helix 9+10 (50) greatly reduce cholesterol efflux and a synthetic 9/10 helix mediates high affinity cholesterol efflux (51). Yet in our studies, ONE modification of HDL did not affect its macrophage cholesterol efflux capacity. As such, our study extends the findings of Shao et al. showing that individual lipid aldehydes varied in their ability to impair specific HDL functions (13) and suggests that specific structural features of various Lys226 adducts determine if there is an effect on cholesterol efflux. Elucidating the specific structural features that account for these differences will require additional studies. One important difference between the lysine adducts of isolevuglandin and ONE may be that the isolevuglandin adducts include a negatively charged carboxylate. Isolevuglandins could also potentially target different lysine residues.

We previously demonstrated the potential of using dicarbonyl scavengers with a 2-aminomethylphenol moiety (e.g. PPM) to block the ability of isolevuglandins to modifying HDL and thus preserve its function. Measurements of the second-order rate constants for the reaction of various scavengers with ONE in vitro showed that PPM is much more reactive than SAM and EtSAM but only slightly more reactive than ClSAM (33). This study did not test the ability of the scavengers to react with ONE in biological systems. We found that PPM is effective at preventing ONE-induced protein crosslinking in HDL, while SAM-based scavengers show little efficacy. In addition, PPM was also able to prevent ONE-induced HDL dysfunction in an *in vitro* model of inflammation. These results combined with our previous study (28) suggest that PPM may have potential use as a therapeutic to protect HDL in vivo.

In conclusion, we show that ONE modifies HDL in humans with FH, who have severe hypercholesterolemia and atherosclerotic disease, and exerts important biological effects; yet, ONE differs significantly in its effects from isolevuglandins, which are similar in terms of reactivity and ability to crosslink proteins. Importantly, ONE crosslinking of HDL proteins does not alter the size or distribution of HDL particles. ONE modification reduces apoA-I exchange but does not alter macrophage cholesterol efflux. ONE modification blocks HDL inhibition of LPS induced macrophage activation, but does not further potentiate cytokine expression. It will be of interest for future studies to determine whether ONE impairs other functions of HDL and its relevance in LDL modifications in atherosclerosis.

## EXPERIMENTAL PROCEDURES

### Materials

Reagents for SDS-PAGE and immunoblotting were from Novex by Life Technologies (Carlsbad, CA). ApoA-I mouse/human (5F4) monoclonal antibody was purchased from Cell Signaling Technology (Danvers, MA). ApoA-II human (EPR2913) monoclonal antibody was purchased from Abcam (Cambridge, MA). OPA reagent was purchased from Thermo Scientific (Rockford, IL). Materials used for cell culture were from Gibco by Life Technologies (Grand Island, NY). [1,2-^3^H(N)]-cholesterol was purchased from Perkin-Elmer Life Sciences. eBioscience LPS solution was purchased from Thermo Fisher Scientific (Waltham, MA). Acetylated LDL derived from human plasma was purchased from Alfa Aesar (Haverhill, MA) RNEasy Mini kit was purchased from Qiagen (Hilden, Germany). iQ SYBR Green Supermix and iScript cDNA Synthesis kit were purchased from Bio-Rad Laboratories (Hercules, CA).

### HDL from FH patients and healthy controls

Ethylenediaminetetraacetic acid plasma was isolated from the blood of FH patients (n=7). Control plasma was isolated from blood of healthy volunteers (n=8). HDL was isolated by FPLC. The study was approved by the Vanderbilt University Institutional Review Board (IRB), and all participants gave their written informed consent. The human blood from FH patients and healthy controls were obtained using an IRB approved protocol.

### Animals

All procedures were approved by the Vanderbilt University Institutional Animal Care and Use Committee (IACUC). Breeding pairs of homozygous apoE^-/-^ mice (C57BL/6J background, strain 002052) aged 12 weeks were purchased from Jackson Laboratories (Bar Harbor, ME). The animals were acclimated and housed in the Vanderbilt University animal facility in a 12-hour light/12-hour dark cycle, were maintained on standard rodent chow (LabDiet 5001), and were given free access to water. Progeny of the breeding pairs were at least 8 weeks of age before harvest of macrophages (described below).

### Chemical Synthesis of ONE and scavengers

ONE was synthesized as previously reported (52) and prepared as a 1M solution in acetonitrile. ONE was diluted as a 10 mM stock solutions in DMSO in aliquots and stored in −80°C until use. Acetic acid salts of salicylamine (SAM), 5-chlorosalicylamine (Cl-SAM), 5-fluoro-salicylamine (F-SAM), and the hydrochloride salt of 5’-O-pentylpyridoxamine (PPM) were synthesized as described (53,54). Working solutions were prepared fresh before each assay and diluted in water to appropriate concentrations.

### Measurement of ONE

Quantitation of lysine modification of HDL by ONE was performed by subjecting an aliquot of HDL to proteolysis with pronase and aminopeptidase M, and then measuring the amount of ONE-ketoamide by stable isotope dilution LC/MS/MS as previously described (55). The internal standard for ONE-ketoamide was synthesized by coupling of 4-oxononanoic acid and ^13^C6-Lys. The N-hydroxysuccinimide ester of 4-oxononanoic acid was coupled to Fmoc-protected ^13^C6-Lys under basic conditions. The product was deprotected with piperidine and purified by flash chromatography on silica gel.

### ONE modification of HDL and the use of scavengers

HDL was exposed to various concentrations of ONE at 37°C overnight to guarantee a complete reaction to form a stable end product. Control HDL was treated similarly in the absence of dicarbonyls. For experiments involving the use of scavengers, scavengers solubilized in water were incubated with HDL for 30 min at 37 °C before the addition of ONE.

### Characterization of apolipoprotein crosslinking of modified HDL

HDL apolipoprotein crosslinking was assessed by SDS-PAGE performed under reducing conditions with Invitrogen’s gel electrophoresis and transfer system. 4-20%Tris gradient gels were used. Western blot analyses were carried out using polyclonal antibodies specific for human apoA-I, apoA-II, and apoA-IV.

### HDL-ApoA-I exchange

HDL samples prepared by adding 15 μL 3 mg/mL spin-labeled apoA-I probe to 45 μL 1 mg/mL HDL and drawn into an EPR-compatible borosilicate capillary tube (VWR) (32). EPR measurements were performed using a Bruker eScan EPR spectrometer outfitted with temperature controller (Noxygen). Samples were incubated for 15 minutes at 37 °C and then scanned at 37 °C. The peak amplitude of the nitroxide signal from the apoA-I probe in the sample (3462-3470 Gauss) was compared to the peak amplitude of a proprietary internal standard (3507-3515 Gauss) provided by Bruker. The internal standard is contained within the eScan spectrometer cavity and does not contact the sample. Since the y-axis of the EPR spectrometer is measured in arbitrary units, measuring the sample against a fixed internal standard facilitates normalization of the response. HDL apoA-I exchange (HAE) activity represents the sample : internal standard signal ratio at 37°C. The maximal % HAE activity was calculated by comparing HAE activity to a standard curve ranging in the degree of probe lipid-associated signal. Experiments were repeated two separate times. All samples were read in triplicate and averaged.

### Cell culture

Male and female apoE^-/-^ mice (C57/BL genetic background) were injected intraperitoneally with 3% thioglycolate and the macrophages were harvested by peritoneal lavage after four days. Cells were maintained in 24-well plates in DMEM with 10% (v/v) fetal bovine serum and penicillin-streptomycin at 100 units/mL and 100 μg/mL respectively.

### Cholesterol efflux

Efflux was assessed by the isotopic method (56). Loading medium was prepared to consist of DMEM containing 100 μg/ml acetylated LDL with 6 μCi ^3^H-cholesterol/ml. After equilibration for 30 minutes at 37°C, loading medium was added to cells for 48 h. After 48h, the cells were incubated for 1h with DMEM containing 0.1% bovine serum albumin so that surface-bound acetylated LDL was internalized and processed. Cells were washed and incubated with efflux medium, which contained DMEM with 35 μg/mL HDL samples. After 24h incubation, supernatants were collected, vacuum filtered, and prepared for β-scintillation counting.

### Macrophage inflammation

Cells derived from female mice were incubated overnight in DMEM containing 0.5% FBS and 1% penicillin-streptomycin. The cells were washed two times with HBSS and then incubated for 4h with DMEM alone or containing 100 ng/ml LPS with or without the HDL preparations (50 μg/ml). The cells were lysed, mRNA harvested, and the cDNA synthesized. qPCR was performed with the following primer pairs: *Tnf* forward (5’-CCATTCCTGAGTTCTGCAAAG-3’); *Tnf* reverse (5’-GCAAATATAAATAGAGGGGGGC-3’); *Il-1β* forward (5’-TCCAGGATGAGGACATGAGCA-3’); *Il-1β* reverse (5’-GAACGTCACACACCAGCA-3’); *Il-6* forward (5’- TAGTCCTTCCTACCCCAATTTCC-3’); *Il-6* reverse (5’-TTGGTCCTTAGCCACTCCTTCC-3’).

### Sample Digestion, Preparation, and LC-MS-MS Analysis of ONE-modified Peptides

Reconstituted HDL synthesized by the cholate dialysis method (57) from recombinant human ApoA-I, phosphatidylcholine, free cholesterol. Particle size homogeneity was checked by native gel electrophoresis. HDL was then dialyzed into PBS before modifying with 3 eq. ONE at 37°C overnight. HDL samples were dialyzed into 50 mM ammonium bicarbonate buffer. In the Davidson lab, the particles were lyophilized to dryness then the lipids were extracted in 1 ml ice cold 2:1 (v:v) chloroform/methanol (58) and the precipitated protein was resolubilized in 80% ammonium bicarbonate buffer and 20% methanol. Samples were reduced by addition of DTT to a final concentration of 10 mM and incubation for 30 min at 42°C. Reduced protein was carbamidomethylated with iodoacetamide at a final concentration of 40 mM and incubated in the dark at RT for 30 min. 20 μg of each sample was digested by adding 1 μg of sequencing grade trypsin (Promega) for 16 h at 37°C. An additional 1 μg of trypsin was added the following day and incubated for an additional 2 h at 37°C. Samples were lyophilized to dryness using a speedvac and stored at −20°C until ready for MS analysis.

Dried peptides were reconstituted in 15 μl of 0.1% formic acid in water and 5 μl of sample was applied to an ACQUITY UPLC C18 reverse phase column (Waters) maintained at 40°C using an Infinity 1290 autosampler and HPLC (Agilent). Peptides were eluted at 0.1 ml/min using a varying mobile phase gradient from 95% phase A (FA/H_2_O 0.1/99.9, v/v) to 32% phase B (FA/ACN 0.1/99.9 v/v) for 120 min followed by 32% B to 50% B for 2 min. Column cleaning was performed by varying the mobile phase gradient to 90% B for 10 min and the column was re-equilibrated at 95% A for 10 min. Peptides were introduced to the mass spectrometer using a Jet Stream source (Agilent) as previously described (59). Spectra was acquired using an iFunnel Q-TOF (Agilent) operating in positive ion mode. Precursors were limited to acquisition of ions with a charge states of 2+ and 3+ and required a minimum of 1500 counts. Each cycle acquired the 20 most intense precursors which were fragmented with a variable collision energy (CE) dependent on the precursor mass-to-charge (m/z) ratio: CE = k* (m/z) +b, with a slope (k) of 3 and an offset (b) of 2 for 2+ ions and −2 for 3+ ions. MS/MS spectra were acquired until at least 45,000 total counts were collected or a maximum accumulation time of 0.33 s. MGF files were generated using MassHunter Qualitative Analysis Software (v B.07.00, Agilent). MS/MS peaks were limited to the top 150 peaks by height and precursors were limited to a maximum assigned charge state of 3+. MS/MS data was analyzed by MassMatrix Suite 3.10 (www.MassMatrix.bio) using settings of peptide mass tolerance of 10 ppm, peptide length of 3-40 amino acids, 1 PTM per peptide, 2 missed tryptic cleavages, and minimum pp score of 5.0. Mass additions for a ONE-ketoamide (C9H14O2, 154.099 Da) or a ONE-Michael adduct (C9H14O2, 156.115) were searched for both Lys and His residues.

## Abbreviations

ONE: 4-oxo-2-nonenal
HNE: 4-hydroxynonenal
HDL: high-density lipoprotein
FH: familial hypercholesterolemia
PPM: pentylpyridoxamine
LDL: low-density lipoprotein
LCAT: lecithin-cholesterol acyltransferase
HPNE: 4-hydroperoxy-2-nonenal
FPLC: fast protein liquid chromatography
EPR: electron paramagnetic resonance
F-SAM: fluoro-salicylamine
Cl-SAM: chloro-salicylamine
MDA: malondialdehyde
HAE: HDL-apoA-I exchange

## Acknowledgements

We acknowledge the Lipoprotein Analysis and HDL Function Core for providing the purified human HDL. This work was supported by National Institutes of Health grants HL116263 (MFL & SSD), HL138745 (LSM). The content is solely the responsibility of the authors and does not necessarily represent the official views of the National Institutes of Health.

## Author contributions

LSM, PGY, MFL, SSD assisted in study concept and design of the studies. SM collected and interpreted experimental data involving HDL crosslinking, biological functions, and scavenging effect. KT synthesized the ONE ketoamide internal standard. VY performed the mass spectrometry analyses for human samples and provided technical support. TP assisted in experimental data collection and analysis involving western blotting. MSB collected and interpreted data related to HAE experiments. AV synthesized ONE and dicarbonyl scavengers. JTM, JM, and WSD collected data, performed the mass spectrometry analysis for amino acid target of ONE and prepared figures representing the results. LSM prepared the figures, drafted and revised the manuscript. MSB, AV, WSD, PGY, MFL, and SSD assisted in analysis and interpretation of data, and provided critical reviews of the manuscript. MFL and SSD obtained project funding, provided technical and material support, supervised all aspects of the study, design, and execution. All authors reviewed the results and approved the final version of the manuscript.

---

**Supplementary Figure 1.**
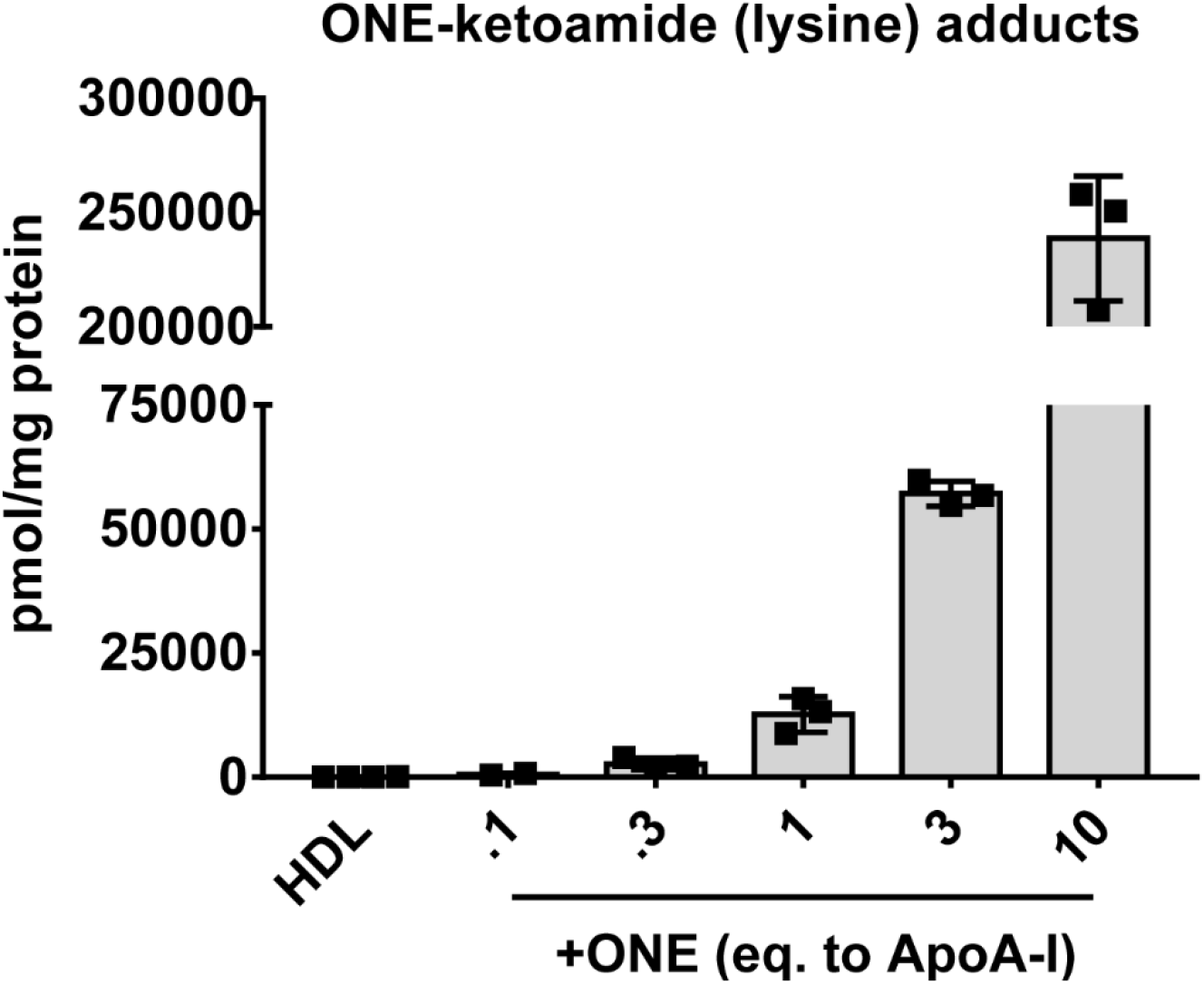
Dose-response of ONE modification to generate ONE-lysine ketoamide adducts in HDL. Control HDL isolated from healthy control volunteers was purified using density gradient ultracentrifugation and was subjected to ONE modification in aqueous conditions for 37°C overnight. Levels of ONE-protein adducts were determined by LC/MS/MS. Results from three individual experiments are plotted as mean ± SD.

